# Evolutionary and structural basis of SLAM utilization in morbilliviruses – Its implications for host range and cross-species transmission

**DOI:** 10.1101/2025.02.18.638793

**Authors:** Ayumu Hyodo, Fumio Seki, Kento Fukuda, Kaede Tashiro, Yuki Kitai, Yukiko Akahori, Hideko Watabe, Hiroshi Katoh, Rikuto Osaki, Daisuke Takaya, Norihito Kawashita, Hideo Fukuhara, Satoshi Ikegame, Tomoki Yoshikawa, Park Eunsil, Shigeru Morikawa, Ryoji Yamaguchi, Benhur Lee, Katsumi Maenaka, Tsuyoshi Shirai, Kaori Fukuzawa, Shigenori Tanaka, Makoto Takeda

**Author notes:** These authors contributed equally to this work (Ayumu Hyodo, Fumio Seki).

## Abstract

Morbilliviruses, including measles virus (MV), canine distemper virus (CDV), and cetacean morbillivirus, pose a significant threat to humans and animals. While the host range of morbilliviruses is generally well-defined, severe cross-species transmission events also have been reported. Their entry into immune cells, the primary targets of morbilliviruses, relies on the signaling lymphocytic activation molecule (SLAM), a receptor whose species-specific variations influence viral host range. However, the extent to which SLAM diversity is a barrier to cross-species transmission remains poorly understood. In this study, we systematically investigated SLAM-mediated host specificity. We found that most morbilliviruses efficiently utilize SLAM from multiple host species, except for human SLAM. Among the morbilliviruses tested, only MV efficiently utilized human SLAM.

Bats are natural reservoirs for many zoonotic viruses. Recent discoveries of novel morbilliviruses in bats provide new insights into morbillivirus evolution. Bats may have played a significant role in morbillivirus evolution because bat (*Myotis*) SLAM also functioned as an efficient receptor for multiple morbilliviruses. Unlike other morbilliviruses, MV utilized *Myotis* bat SLAM inefficiently. We conducted an MV adaptation experiment with *Myotis* bat SLAM to better understand SLAM recognition by morbilliviruses. MV readily adapted to utilize *Myotis* bat SLAM by acquiring a single N187Y mutation in its hemagglutinin protein, and computational structural modeling and fragment molecular orbital calculations with molecular dynamics simulations revealed key interaction changes that facilitated MV’s adaptation to *Myotis* bat SLAM. These findings highlight the adaptability of morbilliviruses in utilizing diverse animal SLAMs. Notably, hypothetical ancestral SLAMs reconstructed in this study acted as universal receptors for all morbilliviruses. These results reinforced that morbillivirus receptor usage is primarily supported by evolutionarily conserved structural features of SLAM, highlighting a molecular basis that enables morbilliviruses to rapidly adapt to diverse animal SLAMs.

**Author Summary:** Our study explores how viruses in the genus morbillivirus, such as measles virus, canine distemper virus, and rinderpest virus, which are notorious for deadly outbreaks in humans and animals, can jump between species. The host range of morbilliviruses is significantly influenced by a receptor molecule on cell surfaces known as the signaling lymphocytic activating molecule (SLAM). By examining SLAMs from various animals, including humans, dolphins, dogs, seals, and bats, we observed how these viruses can infect or adapt to infect different hosts. We found that in some cases slight differences in SLAM may act as initial barriers to cross-species transmission. However, these viruses rapidly overcome such barriers, showing a remarkable ability to adapt. Our research highlights the importance of monitoring these viruses to predict and prevent potential cross-species infections, which is crucial for protecting public health and animal welfare over time.

## Introduction

Morbilliviruses, a genus within the *Paramyxovirus* family, are major pathogens with significant medical and veterinary importance. They pose significant health and economic burdens due to their ability to cause lethal, widespread outbreaks, leading to high mortality rates and economic losses in livestock industries. Each morbillivirus has a distinct host range: measles virus (MV) infects humans, cetacean morbillivirus (CeMV) targets cetaceans, canine distemper virus (CDV) affects carnivores, phocine distemper virus (PDV) infects seals, rinderpest virus (RPV) historically infected cattle, and peste des petits ruminants virus (PPRV) infects small ruminants such as sheep and goats [1, 2]. The recent discovery of novel morbilliviruses in bats (*Myotis*, *Phyllostomus*, and *Molossus*) [3–5], swine [6], and cats [7] suggests a broader host spectrum than previously recognized [2].

Morbillivirus infection begins when the viral hemagglutinin (H) protein binds to a receptor on the host cell surface. The signaling lymphocytic activation molecule (SLAM), expressed on immune cells, and nectin-4, expressed on epithelial cells, serve as primary receptors [1, 2]. Unlike nectin-4, which is highly conserved across species, SLAM exhibits substantial sequence variability [1] (Supplementary Fig. 1), which may significantly influence the viral host range [3, 8–14].

The ability of viruses to jump between species, known as cross-species transmission, can lead to emerging infectious diseases. Such events typically occur when a virus from an animal host acquires genetic mutations that enhance its ability to infect and replicate in humans. Adaptation to host receptor molecules is often a key determinant in defining viral host range, as seen in multiple emerging viruses [15–17]. While paramyxoviruses, including morbilliviruses, can replicate efficiently in human or primate cell lines expressing the appropriate receptor [3, 18], the extent to which SLAM diversity limits or facilitates cross-species transmission remains unclear.

Notably, CDV has caused fatal outbreaks in non-human primates, raising concerns about the potential for morbilliviruses to affect humans [1, 19, 20]. CeMV has also repeatedly caused interorder transmission to seals [1, 21, 22]. This study systematically examines the molecular basis of SLAM-mediated host specificity in morbilliviruses, revealing that conserved structural features of SLAM are crucial for receptor function. Morbilliviruses can often utilize SLAMs from non-native hosts with similar efficiency. Evolutionary comparisons, site-directed mutagenesis, and computational analysis demonstrate that even a few key amino acid changes in SLAM or the viral H protein can substantially alter receptor compatibility. These findings provide critical insights into the mechanisms governing morbillivirus host range and cross-species transmission risk. Understanding how morbilliviruses adapt to new hosts is essential for assessing their zoonotic potential and developing strategies to prevent future spillover events in humans, livestock, and wildlife.

## Results

### Assessment of SLAM utilization by different morbilliviruses

Morbilliviruses have been identified in diverse host species, including humans, cows, goats, dolphins, bats, dogs, and seals. To evaluate their ability to utilize different SLAM receptors, we generated Vero cells expressing SLAM from these species [14, 23, 24] (Supplementary Fig. 2). *Myotis* bat SLAM’s sequence was used as a bat SLAM sequence (Supplementary Fig. 1) [3]. Since antibodies against non-human SLAMs were unavailable, an N-terminal hemagglutinin (HA) epitope tag was introduced, as previously reported [3, 25] (Supplementary Fig. 3). The engineered cells were infected with various morbilliviruses, and cytopathic effects (CPEs), characterized by multinucleated giant cell (syncytia) formation, were observed. Due to regulatory restrictions, live RPV and PPRV were excluded from these experiments [26, 27]. CeMV (muc strain [23]) and CDV (Ac96I strain [18, 28]) induced syncytia in Vero cells expressing dolphin-, bat-, dog-, and seal-SLAMs within 24 hours post-infection (hpi) (Fig. 1A). PDV (A982 strain [23]) exhibited a similar pattern but required up to 96 hpi (Fig. 1B). MV (IC323-EGFP strain [29]) induced syncytia in dolphin-, dog-, and seal-SLAM cells but not in bat-SLAM cells. Notably, MV was the only morbillivirus capable of inducing syncytia in human-SLAM-expressing cells (Fig. 1). The newly identified *Myotis* bat morbillivirus (MBaMV) [3] exhibited a unique specificity, exclusively utilized bat SLAM and failed to induce syncytia in cells expressing SLAM from other species (Fig. 1)

**Figure 1.**
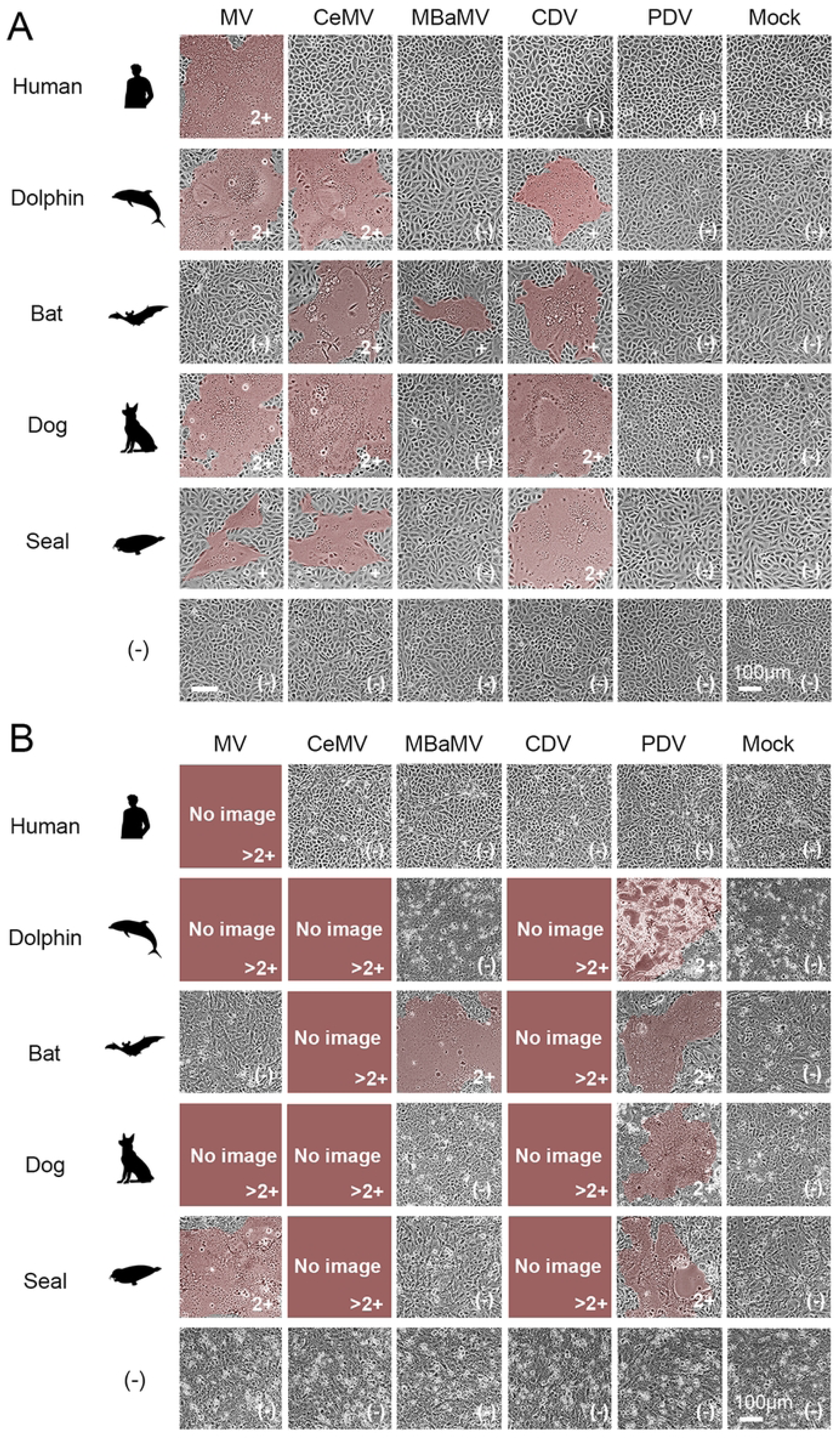
Cytopathic effects induced by morbilliviruses in various SLAM-expressing cells. Vero cells stably expressing SLAM from different animal species (human, dolphin, bat, dog, and seal), along with parental Vero cells (denoted as ’-’), were infected or mock-infected with measles virus (MV), cetacean morbillivirus (CeMV), myotis bat morbillivirus (MBaMV), canine distemper virus (CDV), or phocine distemper virus (PDV) at a multiplicity of infection (MOI) of 0.01. Cytopathic effects (CPEs) were evaluated at 24-hour intervals. (A) CPE observations at 24 hours post-infection (hpi). (B) CPE observations at 96 hours post-infection. CPE scoring: 2+: Large syncytia observed throughout the field of view; +: Few small syncytia detected; (-): No syncytia detected; >2+: Majority of cells detached (no image shown). Syncytial areas are highlighted in brown.

To quantitatively assess viral infectivity, plaque assays were performed in SLAM-expressing Vero cells. MV efficiently formed plaques in human-, dolphin-, dog-, and seal-SLAM cells but showed limited plaque formation in bat-SLAM cells (Fig. 2A). MV produced a higher plaque count in dolphin-, dog-, and seal-SLAM cells than in human-SLAM cells (Fig. 2B), with the largest plaque size observed in dolphin-SLAM cells (Fig. 2A, C). Similarly, CeMV, CDV, and PDV formed plaques in dolphin-, dog-, seal-, and bat-SLAM cells but not in human-SLAM cells (Fig. 2A). In contrast to the broader SLAM utilization seen in these morbilliviruses, MBaMV formed plaques exclusively in bat-SLAM cells (Fig. 2A). Despite its restricted host specificity, bat SLAM efficiently supported plaque formation by CeMV, CDV, and PDV (Fig. 2A), suggesting that bat SLAM retains the key structural motifs necessary for morbillivirus receptor function.

**Figure 2.**
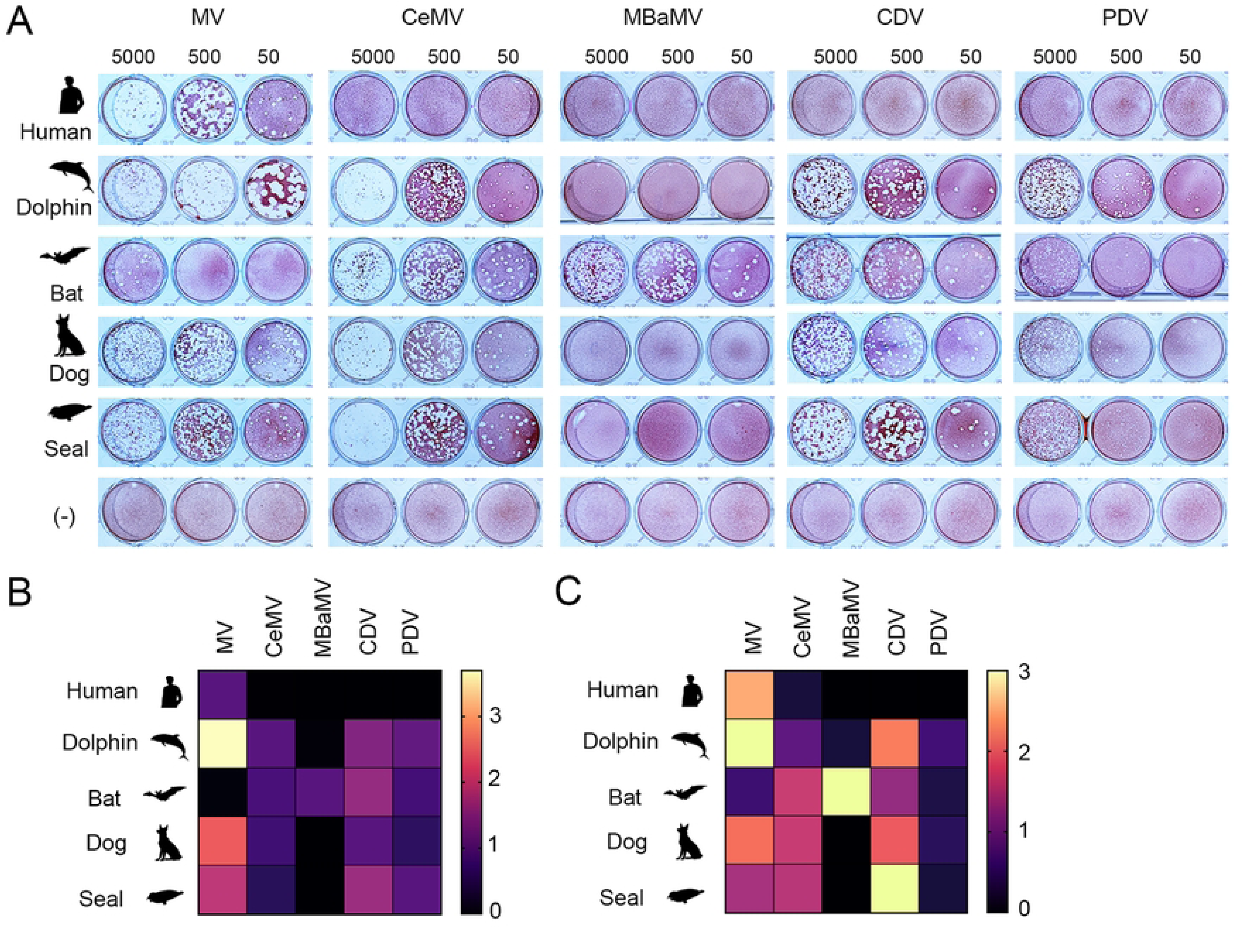
Plaque formation by morbilliviruses in various SLAM-expressing cells. (A) Monolayers of Vero cells stably expressing SLAM from different animal species (human, dolphin, bat, dog, and seal), along with parental Vero cells (-), were infected with different titers (5000, 500, and 50 plaque-forming units [PFUs]) of each morbillivirus (MV, CeMV, MBaMV, CDV, and PDV) and cultured for four days in culture media containing 1% methylcellulose. Cells were stained with neutral red to visualize plaques. The experiment was triplicated, and representative images are shown. (B) Number of plaques formed by each morbillivirus. The mean number of plaques from triplicate analyses in Vero cells expressing the natural host SLAM for each morbillivirus was set to 1, and plaque numbers (means from triplicate analyses) in other cells were displayed in a heatmap. (C) Plaque size (means from triplicate analyses) for each morbillivirus was displayed in a heatmap.

### Evaluation of SLAM utilization by morbillivirus H protein in a fusion assay

The infection and plaque assays using live viruses and SLAM-expressing cells had two major limitations. First, the non-human SLAMs expressed in Vero cells contained an N-terminal hemagglutinin (HA) epitope tag, which could potentially affect receptor function. Second, live RPV and PPRV could not be included due to international regulations [26, 27]. To overcome these limitations, we conducted a dual-split protein (DSP)-based fusion assay, which allowed us to assess SLAM utilization by morbillivirus H and F proteins without using live viruses. Expression plasmids encoding the H and F proteins from various morbilliviruses, including RPV (KabeteO strain [30]) and PPRV (Ghana/NK1/2010 strain [31]), were paired with different animal SLAMs, including cow and sheep SLAMs. Unlike those in Vero cells, these SLAMs had an authentic, unmodified N-terminus. The results from the DSP fusion assay closely mirrored those from the infection assays, confirming the validity of this approach. Cow and sheep SLAMs functioned as receptors for MV, PPRV, CeMV, CDV, and PDV, similar to their respective host SLAMs (Fig. 3). Human SLAM did not support PPRV fusion, reinforcing its restricted usage by non-human morbilliviruses. The DSP assay could not fully assess RPV SLAM usage because the RPV H and F proteins facilitated fusion even in the absence of SLAM (Supplementary Fig. 4). This SLAM-independent fusion is possibly a laboratory-adapted phenotype of RPV, though the KabeteO strain is generally considered to retain wild-type characteristics [30].

**Figure 3.**
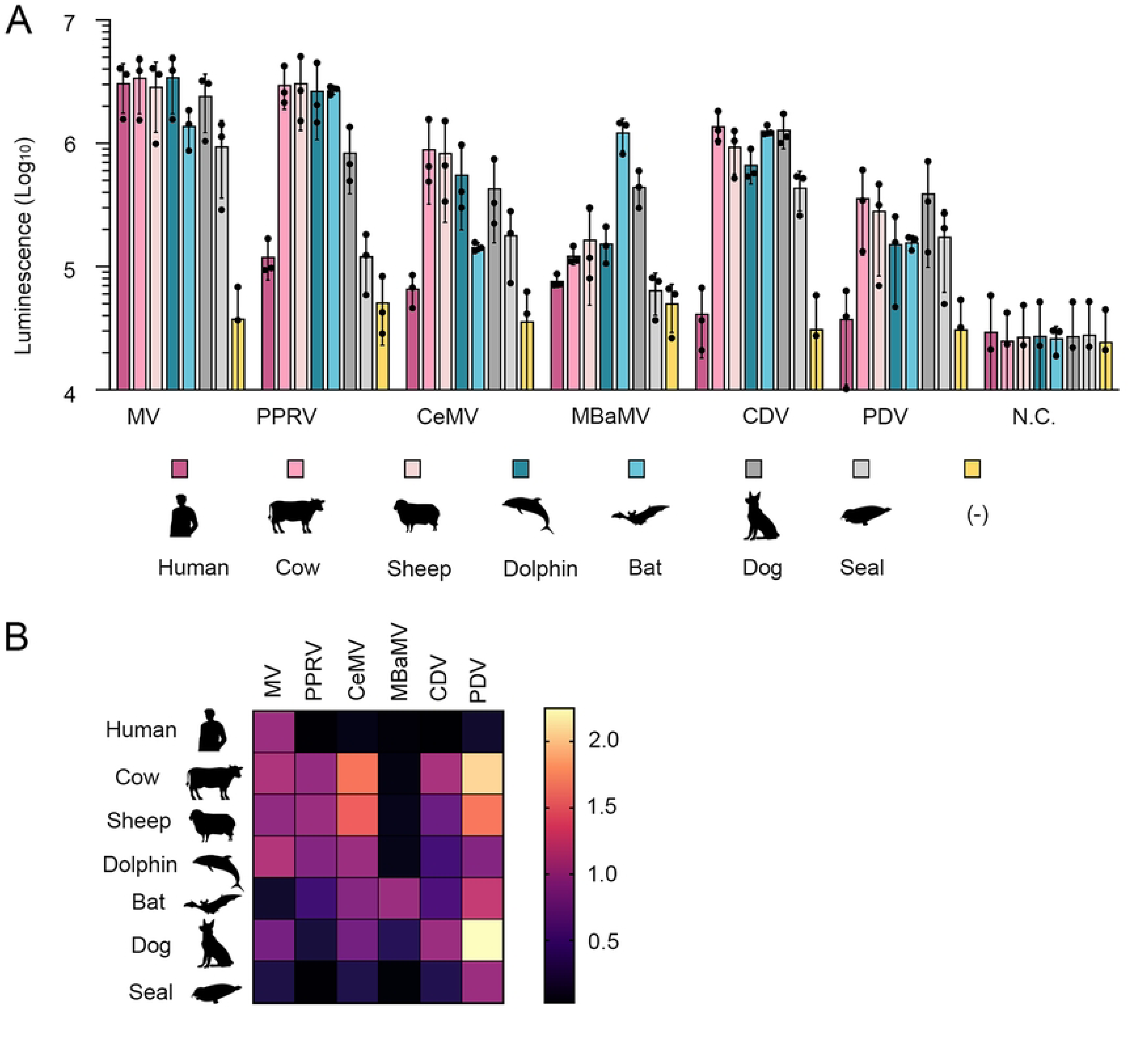
Quantitative cell fusion assay. 293CD4/DSP_1-7_ cells transfected with plasmids encoding different animal SLAMs (human, cow, sheep, dolphin, bat, dog, and seal) and 293FT/DSP_8-11_ cells transfected with both H protein- and F protein-encoding plasmids of each morbillivirus (MV, PPRV, CeMV, MBaMV, CDV, and PDV) were mixed at a 1:1 ratio. *Renilla* luciferase activity was measured three days after transfection. 293FT/DSP_8-11_ cells transfected with the MV F protein-encoding plasmid alone served as a negative control (N.C.). (A) Bar chart displaying raw measured values. Means and standard deviations were calculated from triplicate analyses. (B) Heatmap displaying relative values for each measurement. Values were normalized to 1 when the SLAM of the original host species was used.

### Rapid adaptation of MV to utilize bat SLAM

Unlike other morbilliviruses, MV exhibited poor efficiency in utilizing bat SLAM. We conducted an MV adaptation experiment with bat SLAM to better understand SLAM recognition by morbilliviruses. After several passages in bat-SLAM cells, MV adapted to efficiently replicate and induce syncytia in bat-SLAM cells. Plaque cloning and subsequent assays demonstrated that the bat SLAM-adapted MV produced clear plaques in bat-SLAM cells, with plaque numbers comparable to those in human-SLAM cells (Fig. 4A). Additionally, the adapted MV induced syncytia in bat-SLAM cells, a feature absent in the parental strain but similar to that observed in human-SLAM cells (Fig. 4B). These results indicate that MV readily acquired the ability to efficiently utilize bat SLAM while retaining its capacity to use human SLAM. Sequence analysis revealed a single nonsynonymous mutation (N187Y) in the H gene, where asparagine at position 187 was replaced by tyrosine (Supplementary Fig. 5). To assess its functional impact, the N187Y mutation was introduced into an MV H protein-expression plasmid, and its effect on cell fusion was examined. The mutation significantly enhanced the fusogenic capacity of the MV H and F proteins in bat-SLAM cells, confirming that this minimal amino acid change enabled efficient bat SLAM utilization (Fig. 4C). The analyses up to this point was carried out using bat SLAM with a HA tag (Supplementary Fig. 3). To eliminate potential effects from the HA tag, Vero cells expressing bat SLAM with an authentic, unmodified N-terminus were generated. Analyses using these cells confirmed that MV with the N187Y mutation formed large and numerous syncytia, proliferating in bat-SLAM-expressing cells (Supplementary Fig. 6A). Furthermore, expression plasmid-based analyses reaffirmed that the N187Y mutation was solely responsible for this adaptation (Supplementary Fig. 6B).

**Figure 4.**
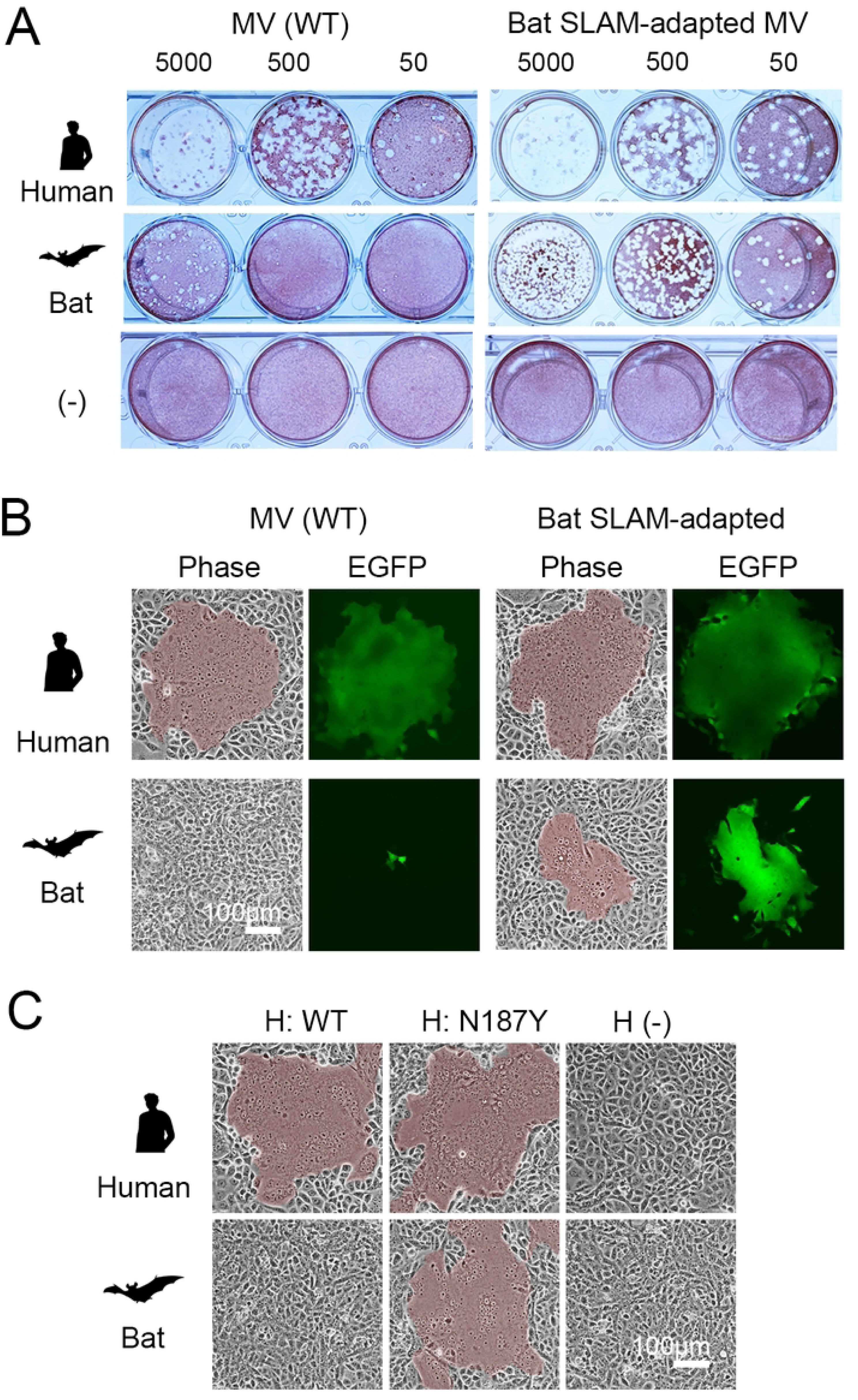
Characterization of bat SLAM-adapted MV. (A) Plaque formation by wild-type (WT) and bat SLAM-adapted MV in Vero cells expressing human SLAM or bat SLAM, as well as parental Vero cells (-). Plaque assays were performed as described in Figure 2. (B) Syncytium formation by EGFP-expressing wild-type and bat SLAM-adapted MV in Vero cells expressing human SLAM or bat SLAM. (C) Syncytium formation induced by H and F protein expression. The MV F protein was co-expressed with either wild-type MV H protein or MV H protein carrying the N187Y mutation in Vero cells expressing human SLAM or bat SLAM. Cells were observed under a microscope one day post-transfection. Syncytial areas are highlighted in brown.

### Computational evaluation of SLAM-H interactions

The adaptation experiment of MV to bat SLAM was conducted using bat SLAM with an N-terminal HA tag (HAtag-batSLAM) (Fig. 5A, Supplementary Fig. 3), leading to the identification of the N187Y mutation. Subsequent biological analyses confirmed that the functional significance of this mutation extends to the native bat SLAM without artificial modifications (Fig. 5A). As an example of morbillivirus adaptation to SLAM receptors from different host species, this study conducted a detailed computational analysis of MV adaptation to utilize bat SLAM. The first calculation procedure was to create a structure of the complex between the MV-H protein and bat SLAM, and perform classical molecular dynamics (MD) calculations thereof. The structure was then extracted from the trajectory of the MD calculations, and the fragment molecular orbital (FMO) calculations were performed on the extracted structure to evaluate the inter-residue interactions.

**Figure 5.**
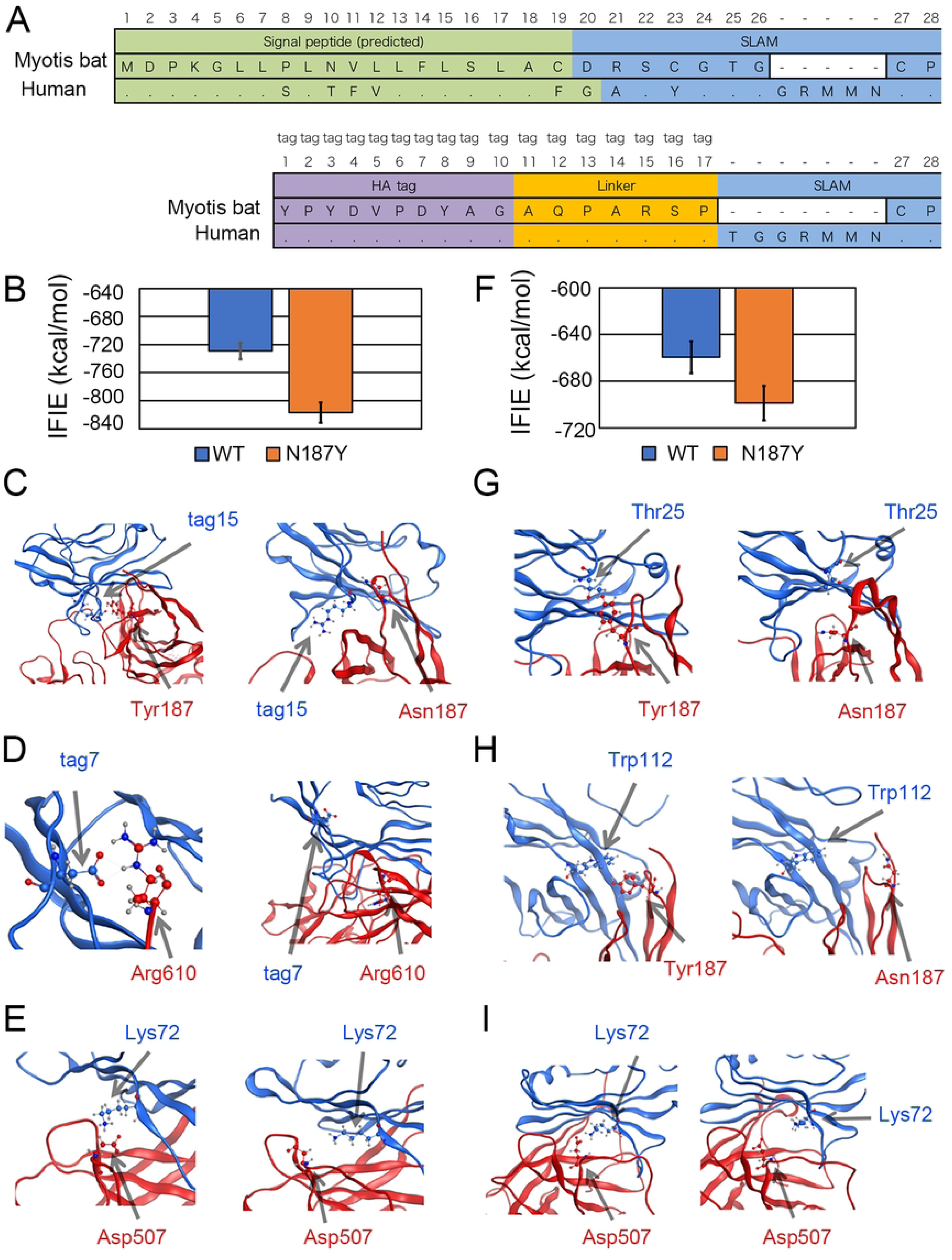
Computational analysis of inter-residue interactions in the complex of MV-H and bat SLAM. (A) Amino acid sequences of bat and human SLAMs in the absence (upper panel) and presence (lower panel) of a tag and linker. Predicted signal peptide sequence, HA tag sequence, SLAM sequence, and linker connecting the HA tag and SLAM sequence are highlighted in green, purple, blue, and orange, respectively. Residues identical to the top sequence are represented as dots, while mismatched residues are shown as their corresponding amino acid letters. Deleted amino acid residues are indicated by hyphens in the alignment analysis. (B) MD trajectory-averaged total IFIE (kcal/mol) between MV-H and bat SLAM with a tag. Blue and orange bars indicate the FMO-calculated results, showing the mean values and standard deviations, for the wild type (WT) and N187Y mutant, respectively. (C) Positional relationship between H-Tyr187 and tag15 (Arg) in bat SLAM with a tag for the N187Y mutant (left) and between H-Asn187 and tag15 (Arg) for the wild type (right). (D) Positional relationship between H-Arg610 and tag7 (Asp) in bat SLAM with a tag for the N187Y mutant (left) and for the wild type (right). (E) Positional relationship between H-Asp507 and Lys72 in bat SLAM with a tag for the N187Y mutant (left) and for the wild type (right). (F) MD trajectory-averaged total IFIE (kcal/mol) between MV-H and bat SLAM without a tag. Blue and orange bars indicate the FMO-calculated results, showing the mean values and standard deviations, for the wild type (WT) and N187Y mutant, respectively. (G) Positional relationship between H-Tyr187 and Thr25 in bat SLAM without a tag for the N187Y mutant (left) and for the wild type (right). (H) Positional relationship between H-Tyr187 and Trp112 in bat SLAM without a tag for the N187Y mutant (left) and for the wild type (right). (I) Positional relationship between H-Asp507 and Lys72 in bat SLAM without a tag for the N187Y mutant (left) and for the wild type (right).

These analyses demonstrated that the N187Y mutation decreased the average inter-fragment interaction energy (IFIE) of the SLAM-H interaction (from −729.0 kcal/mol to −817.0 kcal/mol), enhancing attractive interactions (Fig. 5B). The tyrosine at position 187 (H-Tyr187) formed a hydrogen bond with an arginine (SLAM-tag15) in the HA tag linker (Fig. 5C, Supplementary Fig. 7A). Furthermore, the arginine at position 610 (H-Arg610), near H-Tyr187, strengthened its electrostatic interaction with aspartic acid in the HA tag (SLAM-tag7) (Fig. 5D, Supplementary Fig. 7B, Supplementary Fig. 5). In addition, the aspartic acid at position 507 of the H protein (H-Asp507) significantly interacted with lysine at position 72 of SLAM (SLAM-Lys72) (Fig. 5E, Supplementary Fig. 1). In contrast, in MV-H without the N187Y mutation, SLAM-Lys72 interacted mainly with aspartic acid at position 505 (H-Asp505) instead of H-Asp507 (Supplementary Fig. 7C, D). These two residues (H-Asp505 and H-Asp507) play key roles in MV-H binding to cotton-top tamarin SLAM (PDB ID: 3ALZ) [32] and are highly conserved in morbillivirus H proteins other than MBaMV H protein (Supplementary Fig. 5). This shift in interacting residues, induced by the N187Y mutation in the H protein, further supports its role in enhancing SLAM-H interaction. Next, we analyzed native bat SLAM without an HA tag. Similar to the case of HAtag-batSLAM, the N187Y mutation decreased the average IFIE of the SLAM-H interaction (from −659.5 kcal/mol to −698.8 kcal/mol), strengthening attractive interactions (Fig. 5F). Furthermore, H-Tyr187 formed a new hydrogen bond with threonine near the N-terminus of bat SLAM (SLAM-Thr25) (Fig. 5G, Supplementary Fig. 1, Supplementary Fig. 7E). Additionally, in addition to SLAM-Thr25, H-Tyr187 formed a CH/π interaction with tryptophan at position 112 of SLAM (SLAM-Trp112) (Fig. 5H, Supplementary Fig. 1, Supplementary Fig. 7F). As seen in HAtag-batSLAM, SLAM-Lys72 shifted its interaction from H-Asp505 to H-Asp507 (Fig. 5I, Supplementary Fig. 7G, H), further confirming the role of N187Y in altering SLAM-H binding.

### The receptor function of primate SLAMs for morbilliviruses

Except for MV, all tested morbilliviruses showed limited ability to use human SLAM, suggesting that primate SLAMs may not efficiently function as receptors for these viruses. However, previous studies demonstrated that CDV strains isolated from monkeys (CYN07 [20] and Monkey-BJ01 [33]) can utilize macaque SLAM. To further investigate the receptor function of primate SLAMs, we analyzed the ability of various CDV strains [28, 34–38] to use macaque SLAM and assessed whether this property is restricted to monkey-derived isolates or extends to other CDV strains. Comparative analyses using cells expressing SLAM from macaques, dogs, and humans revealed that most CDV strains isolated from dogs efficiently formed plaques in both macaque- and dog-SLAM cells but not in human-SLAM cells (Supplementary Fig. 8). Infection titers in macaque-SLAM and dog-SLAM cells were comparable, indicating that CDV can efficiently exploit macaque SLAM.

Notably, among dog CDV strains, the P94S strain [28, 34] was uniquely unable to form plaques in macaque-SLAM cells, whereas its closest phylogenetic relative, the Ac96I strain [18], formed plaques efficiently (Fig. 6A; Supplementary Fig. 8). Sequence comparison of the H protein between the P94S and Ac96I strains identified a single amino acid substitution (Y539D; tyrosine to aspartic acid at position 539) (GenBank accession numbers AB212964 and AB753775, respectively) (Supplementary Fig. 5). To evaluate its functional impact, we introduced the Y539D mutation into the Ac96I-H protein-expression plasmid and examined its effect in a fusion assay. The mutation did not affect fusion in dog-SLAM cells but impaired it in macaque-SLAM cells (Fig. 6B), confirming that Y539D is a key determinant preventing CDV P94S from using macaque SLAM.

**Figure 6.**
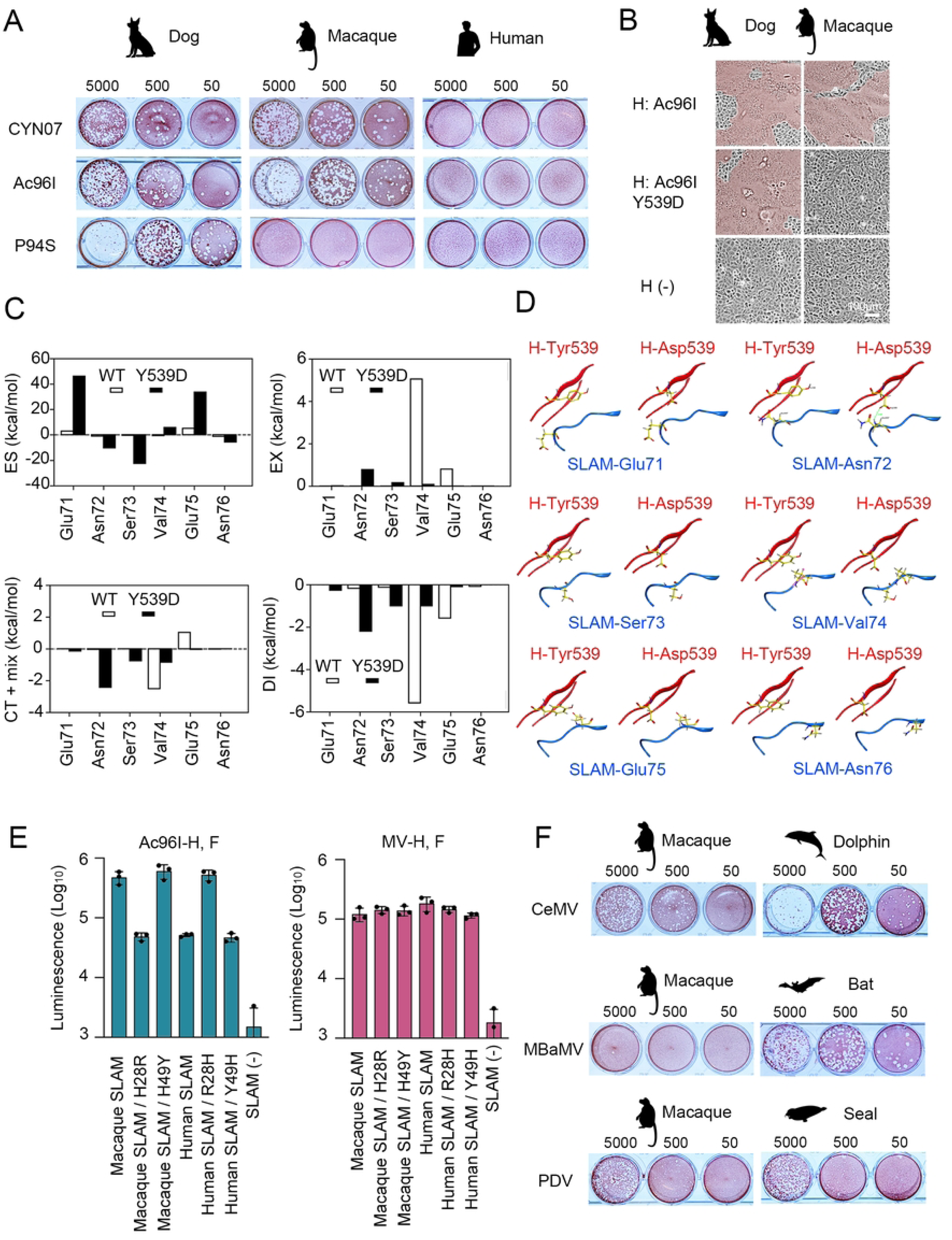
Receptor function of primate SLAMs as morbillivirus receptors. (A) Plaque formation by different CDV strains (CYN07, Ac96I, and P94S) in Vero cells expressing dog SLAM, macaque SLAM, or human SLAM. Plaque assays were performed as described in Figure 2. (B) Syncytium formation induced by H and F protein expression. The F protein of the CDV Ac96I strain was co-expressed with the H protein of the same strain, either with or without the Y539D mutation, in Vero cells expressing dog SLAM or macaque SLAM. Cells were observed under a microscope one day post-transfection. Syncytial areas are highlighted in brown. (C, D) Distribution of interaction energies of each PIEDA term and these interaction pairs in the wild type and Y539D mutant. (C) Bar graphs show the PIEDA energies (ES, EX, CT+mix, and DI) for interactions between Glu71–Asn76 on SLAM and either Tyr539 or Asp539 on CDV-H. (D) The protein structures, displayed in ribbon representation, illustrate the interaction pairs shown in the bar graphs for the wild type (left) and the Y539D mutant (right). (E) Quantitative fusion assay. Mixed cultures of CHO/DSP_1-7_ and CHO/DSP_8-11_ cells were transfected with plasmids encoding macaque or human SLAM, either with or without mutations at amino acid positions 28 or 49, along with plasmids encoding the H and F proteins of CDV (Ac96I strain) or MV. *Renilla* luciferase activity was measured one day post-transfection. Means and standard deviations were calculated from triplicate analyses. (F) Plaque formation by CeMV, MBaMV, and PDV in Vero cells expressing macaque SLAM or their respective natural host SLAM. Plaque assays were performed as described in Figure 2.

Our group previously modeled the Ac96I-H complex with macaque SLAM [39]. FMO calculations were performed for two model structures, wild-type and mutant CDV-SLAM, to quantitatively assess the interactions of the mutated residue and its surrounding residues. The FMO calculation results for wild type and Y539D were performed, and their effects are summarized in Fig. 6C, D. Total IFIE corresponding to the binding energy between CDV-H and the SLAM receptor was −869.8 kcal/mol for the wild-type and the mutant was −820.6 kcal/mol, indicating the interaction with wild-type was more stable by −49.2 kcal/mol.

The changes in Pair Interaction Energy Decomposition Analysis (PIEDA) components between the mutated 539th residue and the SLAM’s residues were observed at SLAM-Glu71 (glutamic acid at 71th residue of SLAM), -Asn72, -Ser73, -Val74, -Glu75 and - Asn76. The sum of IFIE for the six residues was 1.7 kcal/mol in the wild type and 39.5 kcal/mol in the mutant type, indicating that the mutant type is 37.8 kcal/mol less energetically favorable. The effect of the mutation on each interaction was evaluated in terms of the PIEDA components. In particular, the interaction energies for the pair containing SLAM-Glu71, -Val74, and -Glu75 were more attractive in the wild type as compared to the Y539D. The negatively charged H-Asp539 experienced strong electrostatic repulsion from the likewise negatively charged residues SLAM-Glu71 (Δ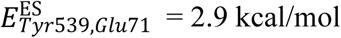 and 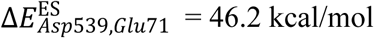) and SLAM-Glu75 (Δ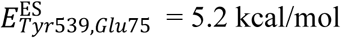 and 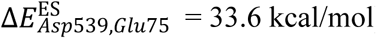), leading to destabilization of the local interaction network. Moreover, the Y539D mutation resulted in the loss of the CH/π interaction between H-Tyr539 and SLAM-Val74 (Δ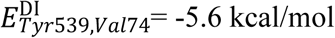 and 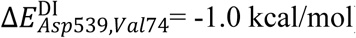), which was present in the wild type. On the other hand, H-Asp539 formed new hydrogen bonds with SLAM-SLAM-Asn73 (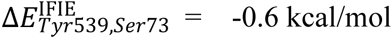 and 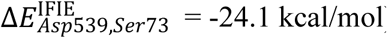) and SLAM-Asn72 (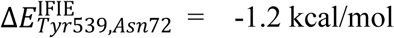 and 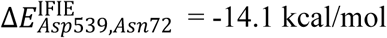) and also strengthened the electrostatic interaction with SLAM-Asn76, contributing to the stabilization of the mutant. Overall, the significant destabilization of IFIE and PIEDA energies caused by the Y539D mutation is consistent with the virological results showing that the Y539D substitution mutant of the CDV-SLAM complex actually lost its binding activity. These FMO calculation results suggest that the Y539D mutation induces repulsion in receptor engagement.

Many CDV strains utilize macaque SLAM but not human SLAM (Fig. 6A; Supplementary Fig. 8). Previous computational studies suggested that a single amino acid difference at position 28 (His28 in macaque SLAM vs. Arg28 in human SLAM) is critical for this specificity [39] (Supplementary Fig. 1). To confirm this, a DSP fusion assay was performed. The results showed that human SLAM with histidine at position 28 (His28) functioned as efficiently as macaque SLAM for CDV entry, while macaque SLAM with arginine at position 28 (Arg28) had reduced receptor functionality (Fig. 6E). Another mutation at position 49, previously identified between human and macaque SLAMs [39], had no effect on receptor function. These findings provide experimental validation of prior in silico predictions [39], confirming that human SLAM cannot serve as a CDV receptor due to the presence of R28.

The receptor function of macaque SLAM for other morbilliviruses (CeMV, MBaMV, and PDV) was also evaluated. CeMV and PDV, but not MBaMV, formed plaques in macaque-SLAM cells at levels comparable to those in their respective host SLAM cells (Fig. 6F). This suggests that macaque SLAM can act as a receptor for multiple morbilliviruses, including CDV, CeMV, and PDV.

### The receptor function of hypothetical ancestral SLAMs for morbilliviruses

Our data indicate that SLAMs from *Carnivora* (dogs and seals), *Cetartiodactyla* (dolphins, cows, and sheep), *Chiroptera* (bats), and *Primates* (macaques) function as receptors for various morbilliviruses, despite considerable differences in their amino acid sequences [1]. We have also prepared mouse SLAM-expressing Vero cells and showed that CeMV, but not other morbilliviruses, used mouse SLAM as efficiently as the host animal (dolphin) SLAM (Supplementary Fig. 9). These findings suggest that the species-specific constraints of SLAM as a morbillivirus receptor are more relaxed than previously assumed, allowing for broader cross-species utilization [3, 8–14]. To explore the evolutionary changes in SLAM and its role as a morbillivirus receptor, we generated and analyzed hypothetical ancestral SLAMs. As the N-terminal five-amino-acid deletion (Supplementary Fig. 1) makes ancestral sequence prediction difficult, bat SLAM was excluded from the analysis. Specifically, we reconstructed ancSLAM8, representing the divergence point between *Primates* (humans) and *Rodentia* (mouse), and ancSLAM9, representing the divergence point between *Cetartiodactyla* (cows, sheep, and dolphins) and *Carnivora* (seals and dogs) (Fig. 7A, B). The DSP fusion assay showed that both ancSLAM8 and ancSLAM9 functioned as receptors for all tested morbilliviruses (MV, CeMV, CDV, PDV, MaBMV, and PPRV) (Fig. 7C). To validate this with live viruses, we generated Vero cells stably expressing ancSLAM8 (Supplementary Fig. 2). These cells formed syncytia (Fig. 7D) and produced plaques with all tested morbilliviruses at levels comparable to their respective host SLAMs (Fig. 7E). These results indicate that the hypothetical ancestral SLAMs functioned as a broadly permissive receptor for morbilliviruses. Furthermore, our data suggest that fundamental structural properties conserved across SLAMs from different animal orders are essential for morbillivirus receptor function.

**Figure 7.**
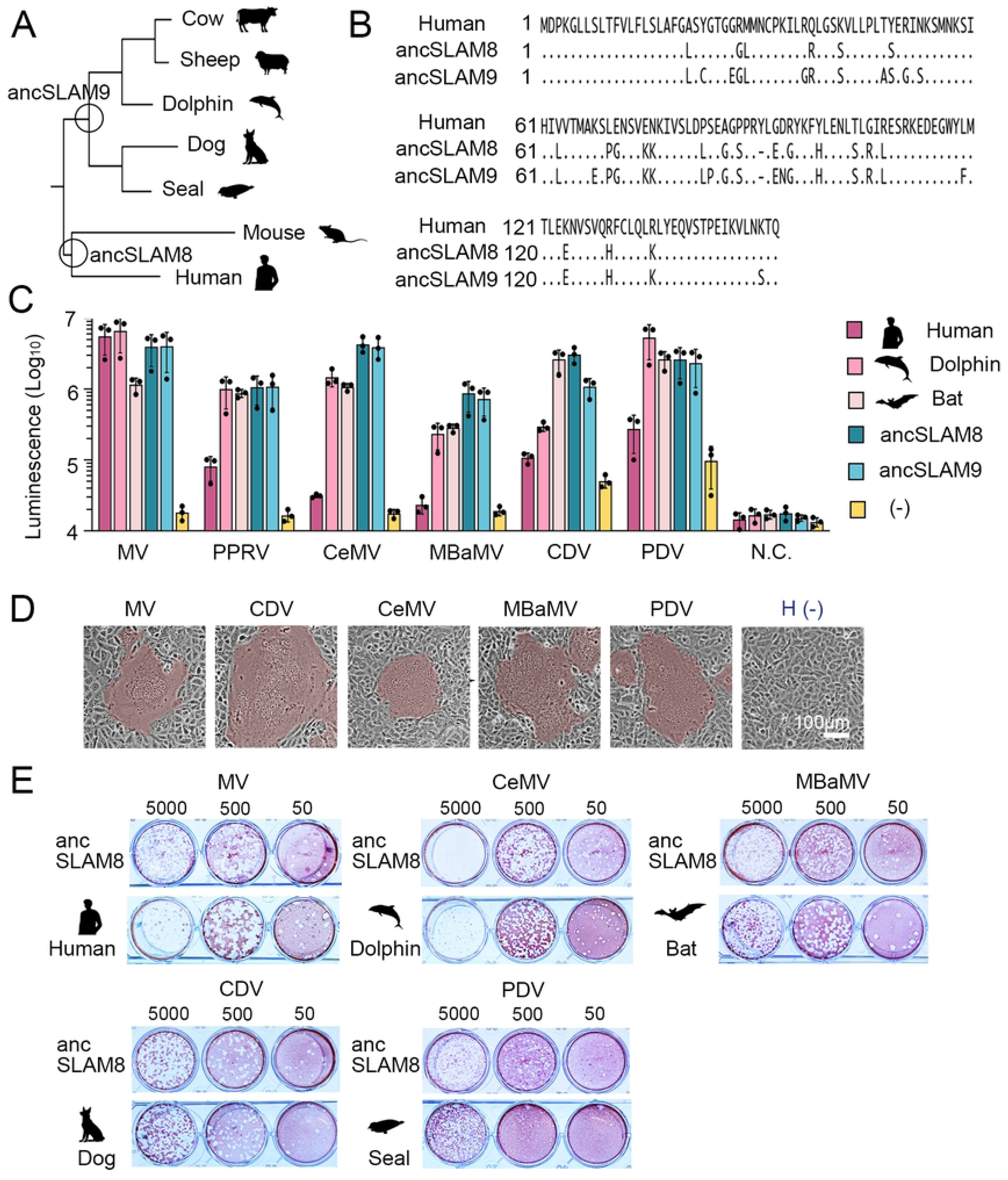
Characterization of hypothetical ancestral SLAMs. (A) Phylogenetic tree constructed with animal SLAMs (cow, sheep, dolphin, dog, seal, mouse, and human). The circles indicate the divergence points where the sequence prediction of ancestral SLAM (ancSLAM8 and ancSLAM9) was performed. (B) Amino acid sequence alignment of the V domains between human SLAM and hypothetical ancestral SLAMs (ancSLAM8 and ancSLAM9). (C) Quantitative cell fusion assay. 293CD4/DSP_1-7_ cells transfected with plasmids encoding different animal SLAMs (human, dolphin, and bat) as well as ancSLAM8 and ancSLAM9, were mixed with 293FT/DSP_8-11_ cells transfected with plasmids encoding the H and F proteins of each morbillivirus. *Renilla* luciferase activity was measured three days post-transfection. 293FT/DSP_8-11_ cells transfected with the MV F protein-encoding plasmid alone were used as a negative control (N.C.). Bar chart shows raw measured values, with means and standard deviations calculated from triplicate analyses. (D) Syncytium formation induced by H and F protein expression. The H and F proteins of each morbillivirus were expressed in Vero cells expressing ancSLAM8 using expression plasmids. Cells were observed under a microscope one day post-transfection. Syncytial areas are highlighted in brown. (E) Plaque formation by MV, CeMV, MBaMV, CDV, and PDV in Vero cells expressing ancSLAM8 or their respective natural host SLAMs. Plaque assays were performed as described in Figure 2.

## Discussion

Morbilliviruses exhibit well-defined host specificity, and species differences in SLAM are important determinants of host range [1, 2]. While the SLAM of the natural host is generally assumed to be the most efficient, our findings suggest that non-host SLAMs can function equally efficiently in many cases [1, 3, 8–14, 25]. Furthermore, hypothetical ancestral SLAMs served as receptors for all tested morbilliviruses. These findings indicate that morbillivirus SLAM utilization is primarily governed by conserved structural features, reducing the impact of sequence variations as a species barrier.

A relatively recent example of morbillivirus evolution is MV, which is believed to have diverged from an ancient bovine morbillivirus around 600 BC [40]. This bovine morbillivirus is also considered the ancestor of RPV. The ability of morbilliviruses to recognize conserved SLAM structures across different mammalian species may have been a key factor in their cross-species transmission and host adaptation. Emerging viruses such as SARS-related coronavirus 2 and ebolavirus have been suggested to originate from bats [15, 17]. Recently, novel morbilliviruses have been identified in various bat species (*Myotis*, *Phyllostomus*, and *Molossus*) [3–5]. Notably, this study showed that *Myotis* bat SLAM efficiently serves as a receptor for various morbilliviruses. Therefore, bats may have played a significant role in morbillivirus evolution.

Unlike other morbilliviruses, MV utilized bat SLAM inefficiently. As an example of morbillivirus adaptation to non-host animal SLAM, we conducted an MV adaptation experiment with bat SLAM and performed a detailed computational analysis of this change. MV successfully adapted to utilize bat SLAM by acquiring a single mutation (N187Y) in the H protein. This mutation enabled the formation of a new hydrogen bond with the N-terminal region of bat SLAM and induced multiple changes in intermolecular interactions. The interaction with the N-terminal region of SLAM is crucial for CDV in utilizing macaque SLAM [39] and for MV in utilizing human SLAM [41]. Our computational analysis data showed that MV even adapted to utilizing the artificially created N-terminal region (HA-tag), providing valuable insights into how the flexibility of protein-protein interactions influences viral host adaptation at the molecular level. These analyses provide insights into molecular interactions between the morbillivirus H protein and the SLAM receptors from both structural and dynamic perspectives.

Our study also provided significant insights into primate SLAMs and morbillivirus infections. Primate SLAMs also retain the conserved structural features essential for morbillivirus receptor function, as macaque SLAM efficiently supported the infection by multiple morbilliviruses. Fatal CDV outbreaks in monkeys [1, 20, 42] have raised serious concerns about the zoonotic potential of non-human morbilliviruses. Our study highlights the unique properties of human SLAM. It functions as a receptor only for MV due to a single SLAM-Arg28 residue. SLAM-Arg28 decreased the interaction energy between SLAM and H protein, when compared with the macaque’s histidine residue at this position (SLAM-His28) [39]. Our data also suggested that SLAM-Arg28 contributed to the incompatibility of human SLAM to the infection with other morbilliviruses including PDV and CeMV. Nonetheless, CDV and PPRV adapt to human SLAM with only one or a few mutations in the H protein [43–45], demonstrating the high adaptation potential of morbilliviruses in utilizing different animal SLAMs.

This study focused solely on SLAM receptors. While receptor usage is a significant factor in the host range, it is not the sole determinant. However, assuming that, even if non-human morbilliviruses could acquire the ability to utilize human SLAM, other restriction host factors protect humans from these viruses is overly optimistic. How can we claim that CDV, which caused mass mortality in monkeys, poses a low risk to humans? Cattle plague (rinderpest) has been one of the most severe infections caused by morbilliviruses, but the pathogen (rinderpest virus) has been eradicated [46]. However, certain morbilliviruses could adapt to cattle and spread, as cow SLAM functions as an efficient receptor for most morbilliviruses. Nectin-4 is another key morbillivirus receptor. Despite its high sequence conservation [1], it may still play a crucial determinant of morbillivirus host range, as it mediates viral shedding from infected animals.

The adaptation to human receptor molecules is one of the crucial steps in the emergence of zoonotic infectious diseases. Indeed, various emerging infectious viruses have exhibited adaptation to human receptor molecules [15–17]. For paramyxoviruses, including morbilliviruses, the receptor specificity represents a critical factor in the cell tropism and host range determination [1, 17, 19, 47]. On the other hand, the roles of other factors remain largely unexplored, except for some involvement of immune-related factors [1, 19, 48]. Generally, non-human animal paramyxoviruses, including morbilliviruses, can replicate in human or primate cell lines if a functional receptor is expressed on the cells [3, 18]. Therefore, the roles of host factors involved in intracellular replication may be less critical than those of receptors for paramyxoviruses.

In conclusion, understanding the contributions of receptor molecules in determining the host range of morbilliviruses is crucial for assessing the basis of cross-species transmission of these viruses and preparing strategies to prevent potential spillover events and outbreaks in humans and livestock. This study provides essential scientific evidence for these efforts.

## Materials and methods

### Cells

Cell lines such as Vero, Vero/hSLAM (human-SLAM), Vero.DogSLAMtag (dog-SLAMs), Vero/dolphin-SLAM (dolphin-SLAM), and Vero/seal-SLAM (seal-SLAM) were previously reported [14, 23, 24]. Vero cells expressing *Myotis* bat SLAM and macaque SLAM were previously described [3, 20]. However, other clones were newly established for this study. Vero cells were transfected with pCAGGS-Igk-HA-bCD150-P2A-Puro [3] and cultured in the presence of 5 µg/ml puromycin (Nacalai Tesque). Puromycin-resistant Vero cell clones were further incubated with 5 µg/ml puromycin. The expression of bat SLAM (bCD150) was confirmed by flow cytometry using an anti-HA tag monoclonal antibody (clone HA11, BioLegend, San Diego, CA, USA) (Supplementary Fig. 2). The established clone was designated as Vero.BatSLAMtag (bat-SLAM) cells. As no specific antibodies are available for dog, dolphin, seal, and bat SLAMs, these SLAMs were tagged with HA epitope at the N-termini, as previously reported [3, 14, 23, 25] (Supplementary Fig. 3). Vero cells were also transfected with pCXN_2_-macSLAM [20] and pCXN_2_-ancSLAM8tag and cultured in the presence of 1.0 mg/ml geneticin (G418; Nacalai Tesque). Geneticin-resistant Vero cell clones were further cultured with 1.0 mg/ml geneticin. Expression of macaque SLAM or ancSLAM8 was verified by flow cytometry using anti-SLAM (clone IPO-3, Kamiya Biomedical) or anti-HA monoclonal (clone HA11) antibodies (Supplementary Fig. 2). The clones established were named Vero/macSLAM-6 (macaque-SLAM) and Vero.ancSLAM8tag cells, respectively. Vero cells expressing bat SLAM with the authentic, unmodified N terminus were also generated. The coding region of Igk-HA-bCD150 in the pCAGGS-Igk-HA-bCD150-P2A-Puro plasmid [3] was replaced with that of bat SLAM with the authentic N terminus, generating pCAGGS-batSLAM-P2A-Puro. Vero cells were transfected with pCAGGS-batSLAM-P2A-Puro and selected in the presence of 10 µg/ml puromycin (Nacalai Tesque). The bulk culture of puromycin-resistant cells, which were predictably expressing bat SLAM, was used for analyses. 293 cells constitutively expressing dual split proteins, namely 293CD4/DSP_1-7_ and 293FT/DSP_8-11_, were reported previously [49]. CHO cells constitutively expressing dual split proteins were developed in this study. CHO/DSP_1–7_ and CHO/DSP_8–11_ cells were generated by transfecting CHO cells with pRL-DSP_1–7_-neo or pRL-DSP_8–11_-neo, respectively, and selected in DMEM supplemented with 7% FBS and 0.5 mg/ml geneticin. All cell lines were maintained in Dulbecco’s Modified Eagle Medium (DMEM) supplemented with 7.5% fetal calf serum (FCS) and antibiotics (100 U/ml penicillin and 0.1 mg/ml streptomycin).

### Viruses

Recombinant wild-type MV expressing enhanced green fluorescent protein (EGFP) has been described previously [29]. Similarly, CeMV muc strain and PDV 982A strain, as well as MBaMV expressing EGFP, have been previously reported [3, 23]. The CDV strains isolated using dog-SLAM cells, including Ac96I, S124C, P94S, MD231, MS232, MSA5, Th12, 007Lm, 009L, 011C, M24Cr, and 55L, were also detailed in previous studies [28, 34–38, 50]. Additionally, the CDV strain isolated from a cynomolgus monkey, designated CYN07-dV, has been previously described [20].

### Plasmid constructions

Mammalian expression plasmids encoding human SLAM (pCA7-hSLAM) and bat SLAM (pCAGGS-Igk-HA-bCD150) were previously described [3, 51]. A mammalian expression plasmid encoding bat SLAM with the authentic N terminus was generated by replacing the coding region of Igk-HA-bCD150 in the pCAGGS-Igk-HA-bCD150-P2A-Puro plasmid [3] with that of bat SLAM with the authentic N terminus, generating pCAGGS-batSLAM-P2A-Puro. Mammalian expression plasmids encoding SLAMs for dolphins, dogs, seals, cows, and sheep were constructed by cloning the coding regions of each respective animal SLAM gene into the pCAGGS vector or its derivatives [52]. The DNA fragments encoding these SLAM genes were chemically synthesized based on the sequences available in GenBank, with accession numbers NM_003037.4 (human), XM_004327846.1 (bottlenose dolphin), AF325357 (dog), AB428368 (spotted seal), XP_014402801.1 (riparian myotis bat), BC114833.1 (cow), and DQ228866.1 (sheep) (Supplementary Fig. 1). Additionally, DNA fragments encoding hypothetical ancestral SLAM sequences (ancSLAM8 and ancSLAM9) were chemically synthesized, and the cDNA of chimeric SLAM genes with the V domain of ancSLAM8 or ancSLAM9 and the C2 domain of human SLAM were inserted into the pCAGGS vector. The chimeric SLAM cDNA with the V domain of ancSLAM8 and the human SLAM C2 domain was also inserted into the pCXN_2_ vector with an N-terminal HA-tag, and the constructed plasmid was named pCXN_2_-ancSLAM8tag.

Additionally, mammalian expression plasmids encoding N-terminally HA-tagged human and macaque SLAM (GenBank accession numbers NM_003037.4 and XM_001117605.3, respectively) were generated using the pCAGGS vector or its derivatives, as previously reported [3, 14, 23, 25] (Supplementary Fig. 2). Site-directed mutagenesis via PCR was employed to introduce amino acid substitutions R28H and Y49H into the HA-tagged human SLAM-expression plasmid, and H28R and H49Y into the HA-tagged macaque SLAM-expression plasmid.

Mammalian expression plasmids for the H and F proteins of MV (wild-type IC-B strain) and the H protein of CDV (Ac96I strain) were previously described [18, 53]. Mammalian expression plasmids for the F protein of CDV (Ac96I strain), and the H and F proteins of PPRV, RPV (KabeteO strain), PDV (982A strain), CeMV (muc strain), and MBaMV were constructed by inserting the coding regions of each protein gene into the pCAGGS vector or its derivatives. These viral gene sequences were also synthesized chemically based on GenBank entries, with accession numbers provided in Supplementary Table 1. Site-directed mutagenesis was used to introduce mutations; N187Y into the MV H protein-expression plasmid and Y539D into the CDV H protein-expression plasmid.

Expression plasmids for the DSP system (pRL-DSP_1–7_ and pRL-DSP_8–11_) were kindly provided by Dr. Z. Matsuda [54, 55]. The region containing the SV40 enhancer/promoter and the coding region of neomycin phosphotransferase was excised from pCI-neo (Promega) and cloned into pRL-DSP_1–7_ and pRL-DSP_8–11_, creating pRL-DSP_1–7_-neo and pRL-DSP_8–11_-neo, respectively.

### Virus infection assay

Sub-confluent monolayers of cells (specific animal SLAM-expressing Vero cells or their parental Vero cells) were infected with morbilliviruses of interest, and the cytopathic effect (CPE) in the cells was observed daily for 4 days.

### Plaque assay

Sub-confluent monolayers of cells (specific animal SLAM-expressing Vero cells or their parental Vero cells) were incubated with tenfold serial dilutions of virus samples for 1 hour. The cells were then overlaid with DMEM supplemented with 1% methylcellulose and 7.5% fetal calf serum (DMEM/MC/FCS). Four days post-infection, DMEM/MC/FCS containing neutral red was added to the cultures. Plaques stained by neutral red were counted the following day.

### Conventional cell fusion assay

Sub-confluent monolayers of cells (specific animal SLAM-expressing Vero cells or their parental Vero cells) cultured in 12-well cluster plates were transfected with plasmids encoding the morbillivirus H and F proteins of interest (0.5 µg per well each). Then, the cells were observed for syncytium formation under a microscope daily for 2 days.

### Quantitative cell fusion assay

293FT/DSP8_-11_ cells, cultured in 6-well plates, were transfected with plasmids encoding specific animal SLAM (1.25 µg per well) using the TransIT-293 transfection reagent. Simultaneously, 293CD4/DSP_1-7_ cells cultured in separate 6-well plates were transfected with plasmids encoding the morbillivirus H and F proteins (2.5 µg and 1.25 µg per well, respectively). Twenty-four hours post-transfection, cells from each culture were resuspended in 3 ml of fresh medium (DMEM/MC/FCS). Then, 92 µl of each resuspended cell suspension was mixed in various combinations and seeded into 96-well plates. After a 15-hour incubation, *Renilla* luciferase activity was quantified using the *Renilla*-Glo Luciferase Assay System (Promega).

For assays using CHO/DSP_1–7_ and CHO/DSP_8–11_ cells, a 1:1 cell mixture was seeded into 24-well plates. These cells were transfected with plasmids encoding the specific animal SLAM receptor (0.125 µg per well) and the morbillivirus H and F proteins of interest (0.125 and 0.25 µg, respectively, per well) using FuGENE HD. *Renilla* luciferase activity was measured the next day using the *Renilla*-Glo Luciferase Assay System (Promega)

### Ancestral SLAM reconstruction

Amino acid sequences of each SLAM were obtained from NCBI with the following accession numbers: human (NP_003028.1), house mouse (AAF22231.1), common bottlenose dolphin (XP_004327894.1), cattle (AAI14834.1), sheep (ABB58749.1), dog (AAK61857.1), and spotted seal (BAH10672.1). Protein sequences were aligned utilizing Clustal Omega [56]. Using the parameters of Jones-Taylor-Thornton (JTT) model, phylogenetic trees were subsequently generated by the neighbor-joining method with MEGA X [57]. Finally, ancestral SLAM sequences were reconstructed using PAML [58], employing the parameters of the Jones-Taylor-Thornton (JTT) model.

### Computational analysis of the CDV-H Y539D mutation by FMO calculations

The input 3D molecular structures for FMO calculations [59, 60] were constructed using MOE [61], employing the wild-type (CDV-H and macaque SLAM complex) model previously developed by our research group [39]. The Y539D mutant structure was constructed from the wild-type model using the Protein Builder module. Generation of hydrogen atom and energy minimization were performed by Structure Preparation module implemented in MOE.

In the energy minimization process, constraint conditions were introduced: atoms within a 4.5 Å radius of the mutated residue (tether = 1.0) and side chain (tether = 0.5). No constrains were introduced for hydrogen atoms. AMBER10:EHT force field was used in all process. FMO calculations were performed on the Fugaku supercomputer, utilizing the ABINIT-MP software (version 1.23) [62]. Fragmentation of proteins was performed in default setting. The IFIE refers to the effective interaction between a pair of specific residues evaluated by the FMO method with inclusion of electron correlation effects by means of Møller-Plesset second-order (MP2) perturbation theory with 6-31G* basis set (FMO-MP2/6-31G*) [63, 64]. IFIE and electrostatic interaction (ES), exchange repulsion (EX), charge transfer with higher-order mixed-term interactions (CT+mix) and dispersion interaction (DI) of PIEDA were employed for the evaluation of the interactions. The interaction energy between CDV-H and the SLAM receptor was assessed using IFIE. PIEDA was utilized to analyze the interactions involving Glu71, Asn72, Ser73, Val74, Glu75, and Asn76, which constitute interaction sites between CDV-H and SLAM.

### Computational analysis of the MV-H N187Y mutation by MD and FMO calculation

The structures of the individual proteins, MV-H (residues 180–610, strain IC-B: NC_001498.1) and bat SLAM (residues 20–139, *Myotis brandtii*: XP_014402801.1) were created using AlphaFold2 [65] based on their respective amino acid sequences. Additionally, the structure of bat SLAM was modeled with an HA tag consisting of 17 residues combined with SLAM residues 27–139 (see Fig. 5A). The individual protein structures were then superimposed using MOE [61] onto the complex structure of MV-H and cotton-top tamarin SLAM (PDB ID: 3ALZ) [32], generating three-dimensional models of the MV-H and bat SLAM complex, both with and without the HA tag. The residue at position 187 in MV-H was subsequently mutated from asparagine (N) to tyrosine (Y) (N187Y) using MOE, resulting in four different MV-H and bat SLAM complex structures: with or without the HA tag on bat SLAM and with or without the N187Y mutation in MV-H. Finally, homology modeling was performed on the N-terminal regions of SLAM in the four complex structures (residues 20–31 for the untagged structure, residues 1–17 of the HA tag, and SLAM residue 27 for the tagged structure; see Fig. 5A) to refine any unnatural structural features. In the procedure described above, AlphaFold2 was executed on Google Colab [66] without a template, and the structure with the highest pLDDT (predicted local distance difference test) score was selected from the five generated models. For structural optimization using MOE, the AMBER10:EHT force field was employed.

MD calculations on the modeled complex structures in a periodic simulation cell of solvated water were executed using GROMACS [67] on the Fugaku supercomputer. Structural sampling was performed to extract snapshots from the simulation trajectory. TIP3P [68] was used as the water force field, and ff14SB [69] was employed as the protein force field. The calculation protocol consisted of structure optimization, heating process, density relaxation, equilibration process, and main calculation, in that order. Temperature and pressure were controlled using the Berendsen thermostat and the C-rescaling method, respectively. Five independent 100 ns MD calculations were performed for structures containing tagged SLAM. Eleven snapshots were extracted from each trajectory at 10 ns intervals, yielding 55 structures. Large fluctuations, primarily associated with SLAM-Asp20, were observed for structures containing untagged SLAM. Therefore, ten independent 100 ns MD calculations were performed to eliminate outlier structures exhibiting such fluctuations. Eleven snapshots were extracted from each trajectory at 10 ns intervals, yielding 110 structures. These structures were then subjected to structural filtering. First, structures in which the nearest neighbor distance exceeded 2.0 Å were excluded. This criterion was applied to the distance between SLAM-Lys72 and its strongest interacting residue, H-Asp505 in the wild type and H-Asp507 in the N187Y mutant, and the distance between SLAM-Glu119 and H-Arg533. Subsequently, 55 structures were selected based on the increasing distance between SLAM-Asp20 and the nearest residue in the MV-H protein.

FMO calculations were conducted on the extracted structure to assess inter-residue interactions. For each of the 55 structures thus obtained and locally optimized by MOE, FMO-MP2/6-31G* calculations were performed in the same way as for the CDV-H Y539D mutation. The overall interaction energy between the MV-H and SLAM proteins was calculated by summing the IFIE values.

## Acknowledgments

We thank Drs. Zene Matsuda and Mizuki Yamamoto for providing the DSP system, and Dr. Yusuke Yanagi for providing various reagents related to SLAM and recombinant MVs. We also thank Drs. Tadashi Maruyama, Yoshiharu Mori, Kazue Ohishi, Hiroaki Tokiwa, and Yuta Yamamoto for their invaluable support and suggestions. We also sincerely thank Kyoryu Pharmaceutical Co., Ltd. for providing CDV strains for this study. This study was performed as part of the activities of the FMO Drug Design Consortium (FMODD) using the Fugaku supercomputer (project ID: hp240162).

## Author Contribution

**Conceptualization** Ayumu Hyodo, Shigenori Tanaka, Makoto Takeda

**Data Curation** Rikuto Osaki, Kaori Fukuzawa, Shigenori Tanaka

**Formal Analysis** Rikuto Osaki, Kaori Fukuzawa, Shigenori Tanaka

**Funding Acquisition** Hiroshi Katoh, Makoto Takeda

**Investigation** Ayumu Hyodo, Fumio Seki, Kento Fukuda, Kaede Tashiro, Yuki Kitai, Yukiko Akahori, Hideko Watabe, Hiroshi Katoh, Rikuto Osaki, Daisuke Takaya, Norihito Kawashita, Hideo Fukuhara, Tomoki Yoshikawa, Park Eunsil, Katsumi Maenaka, Shigenori Tanaka, Makoto Takeda

**Methodology** Ayumu Hyodo, Fumio Seki, Tsuyoshi Shirai, Kaori Fukuzawa, Shigenori Tanaka, Makoto Takeda

**Project Administration** Hiroshi Katoh, Shigenori Tanaka, Makoto Takeda

**Resources** Fumio Seki, Hiroshi Katoh, Satoshi Ikegame, Shigeru Morikawa, Benhur Lee, Kaori Fukuzawa, Shigenori Tanaka, Makoto Takeda

**Software** Kaori Fukuzawa, Shigenori Tanaka

**Supervision** Shigenori Tanaka, Makoto Takeda

**Validation** Ayumu Hyodo, Fumio Seki, Kento Fukuda, Tsuyoshi Shirai, Shigenori Tanaka, Makoto Takeda

**Writing – Original Draft Preparation** Ayumu Hyodo, Shigenori Tanaka, Makoto Takeda

**Writing – Review & Editing** Ayumu Hyodo, Shigenori Tanaka, Makoto Takeda

